# Accurate highly variable gene selection using RECODE in transcriptome data analysis

**DOI:** 10.1101/2025.06.23.661026

**Authors:** Yusuke Imoto

## Abstract

Recent transcriptomics technologies enable gene-expression profiling at single-cell or micrometer-scale spatial resolution, but capture only a small fraction of true RNA molecules, introducing substantial technical noise driven by random sampling. These noise effects distort the earliest analytical steps, dimensionality reduction or highly variable gene (HVG) selection, and their consequences propagate into downstream analyses. The central aim of this study is to address this issue fundamentally by appropriately removing technical noise at its source. Here, I demonstrate that HVG selection based on RECODE, a de-noising method grounded in high-dimensional statistical theory, outperforms widely used approaches for both scRNA-seq and spatial transcriptomics data. RECODE-based HVG selection achieves higher accuracy and robustness, avoids missing values, improves down-stream performance, and provides the fastest runtime and best scalability among noise-reduction methods. These findings show that theory-driven noise removal is essential for recovering true biological signals and establish RECODE as a practical and reliable preprocessing strategy for single-cell analysis.

## Introduction

Recent advances in single-cell RNA sequencing (scRNA-seq) and spatial transcriptomics now enable transcriptome-wide profiling at the resolution of individual cells or even micrometer-scale spatial coordinates. These technologies have provided unprecedented insights into developmental processes, cellular heterogeneity, and disease mechanisms [1]. Despite this progress, scRNA-seq and spatial transcriptomics measurements capture at most about ten percent of the RNA molecules actually present in a cell, generating substantial technical noise, including dropout events [2]. Moreover, because transcriptomic measurements are intrinsically high-dimensional, noise accumulates in downstream computations, leading to the well-known curse of dimensionality, which degrades the reliability of virtually all analytical steps [3]. Without properly addressing these noise-related issues, it becomes difficult to fully extract the biological information contained in single-cell data, hindering tasks such as detecting rare cell populations or characterizing lowly expressed functional genes.

A variety of strategies have been proposed to mitigate noise in single-cell data analysis. One approach is to apply dimensionality reduction methods such as principal component analysis (PCA), but PCA is mathematically known to suffer from the curse of dimensionality and therefore cannot fundamentally resolve the underlying noise problem [4]. Another strategy is noise reduction known as imputation, which attempts to denoise the raw sequencing data. Many imputation methods, including smoothing-based methods (e.g., MAGIC [5]) and model-based approaches (e.g., SAVER [6]), have been shown to improve certain down-stream analyses but often worsen others, a phenomenon known as cyclicity, indicating incomplete noise removal [7, 8]. A third line of approaches involves selecting highly variable genes (HVGs) to reduce the effective dimensionality [9]. However, because technical noise varies substantially across genes, the variability statistics of each gene must be properly adjusted. Existing HVG selection methods address this issue in different ways, such as normalizing dispersions [10], regressing the mean-variance relationship using negative binomial distribution [11, 12, 13], or leveraging zero-count enrichments [14]. While HVG selection mitigates some issues of high dimensionality, it does not remove noise itself and thus does not fully guarantee reliable downstream analyses.

To fully ensure consistent and reliable downstream analysis, one must remove technical noise at its source and recover the underlying true biological signal. To address this challenge, I previously developed RE-CODE (resolution of the curse of dimensionality), a mathematically grounded denoising framework for single-cell omics data [15]. RECODE models the additive technical noise introduced during the measurement process and removes it using eigenvalue modification theory from high-dimensional statistics [16]. The eigenvalues of the RECODE-denoised covariance matrix coincide with the modified eigenvalues, which possess proven asymptotic properties and allow valid variance estimation even when the dimension exceeds the sample size. More recently, the incorporation of eigenvector correction [17] has further improved denoising accuracy in an updated version of the method [18]. Because RECODE performs noise removal based on first-principle statistical theory rather than heuristic smoothing or empirical normalization, it faithfully preserves the true biological structure of single-cell data and provides theoretically supported variance estimates for downstream analyses.

As a consequence, RECODE has been shown to improve the accuracy of a broad range of downstream analyses, including clustering, trajectory inference, and differential expression analysis [19, 20, 21], without exhibiting cyclicity, which commonly affects imputation methods. In practice, the application of RECODE has enabled important biological discoveries, such as identification of functional genes and rare cell types in human and cynomolgus monkey germ cells [20, 22], human neuroblastoma [23], human T cells [24], and mouse intestinal stem cell and chondrocyte lineages [25, 26].

Although RECODE theoretically enhances all downstream analyses, its quantitative evaluation relative to existing methods has remained limited. In this study, I focus specifically on HVG selection, which is the earliest and most influential step in single-cell analysis pipelines, and propose RECODE-based HVG selection. RECODE-based HVG selection is simple and interpretable: after denoising the data with RECODE, I perform total-count normalization and log transformation, standard preprocessing steps in single-cell analysis, and then select genes with the largest variances. I compare RECODE’s HVG selection procedure against widely used existing methods, including SCTransform [11, 27] used in the latest version of Seurat [28], using both scRNA-seq and spatial transcriptomics datasets. I assess the accuracy of HVG selection using known marker genes, evaluate robustness under random subsampling of cells, and quantify improvements in downstream clustering performance. In addition, I examine computational efficiency and scalability to provide practical guidance for users.

Through this comprehensive benchmarking, I demonstrate that RECODE effectively resolves the fundamental noise-related challenges in single-cell data analysis and serves as a reliable and powerful preprocessing method for both scRNA-seq and spatial transcriptomics.

## Materials and Methods

### Overview of RECODE

I provide an overview of the resolution of the curse of dimensionality (RECODE) algorithm. For details, refer to the Supplementary File of Imoto–Nakamura et al. [15] for version 1 and the Methods section of Imoto [18] for version 2.

I first prepare notations and introduce a technical noise in single-cell omics data. Let 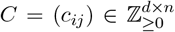 denote the raw count matrix of scRNA-seq data, where *c*_*ij*_ represents the observed RNA (UMI) count for gene *i* in cell *j*. Here, *d* and *n* indicate the number of genes and cells, respectively. Cells with zero total count (∑_*i*_ *c*_*ij*_ = 0) and genes with zero total count (∑_*i*_ *c*_*ij*_ = 0) are excluded from the analysis. I set a total count of cell *j* as *t*_*j*_ := ∑_*i*_ *c*_*ij*_ and a scaled count value for gene *i* in cell *j* as *x*_*ij*_ := *c*_*ij*_*/t*_*j*_. I use super script notation (·)^true^ as the corresponding true data, as 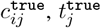, and 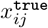. Then, I define technical noise as the difference between true and observed scaled data, i.e., 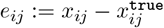.

Modeling the single-cell omics data creation process and analyzing it, I proved the following statistical relationship (Theorem 4.1 in Supplementary File of Imoto–Nakamura et al. [15]):

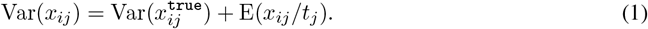

Here, E and Var denote the expectation and variance for gene direction, that is, Var(*x*_*ij*_) shows the variance of gene *i* for data *x*_*ij*_. Further, under the independence of true data and noise, I obtain the variance of noise:

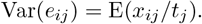

The noise variance Var(*e*_*ij*_) = E(*x*_*ij*_*/t*_*j*_) can be empirically computed as 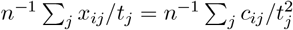. Because Equation (1) leads Var(*x*_*ij*_) ≥ E(*x*_*ij*_*/t*_*j*_) ≥ 0, the observed variance Var(*x*_*ij*_) has a lower bound imposed by the noise variance Var(*e*_*ij*_), regardless of the true signal variance 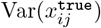 (Figure S1). This proves the existence of lower bound in the distributions of variety statistics, which is discussed in the subsequent section, *Technical noise floor curve disturbs accurate HVG selection*.

Next, I introduce a noise variance-stabilizing normalization (NVSN), which is the first step in the RECODE algorithm, defined as

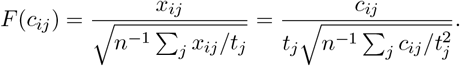

NVSN equalizes the noise variance, letting the lower bounds flat, i.e., Var(*F* (*e*_*ij*_)) = 1 and Var(*F* (*x*_*ij*_)) ≥ 1. which originates from the scRNA-seq measurement process and is mathematically modeled as technical noise. This step is also essential to ensure compatibility with the subsequent eigenvalue and eigenvector correction procedures developed in the field of high-dimensional statistics.

Next, the covariance matrix of the normalized matrix *F* (*C*) = (*F* (*c*_*ij*_)) ∈ ℝ ^*d*×*n*^ is computed as 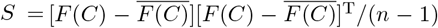, where the notations of overline and superscript T represent a matrix in which the mean vector of each row is replicated across all columns and the matrix transpose, respectively. The eigenvalue equation *Su*_*i*_ = *λ*_*i*_*u*_*i*_ (∥*u*_*i*_∥ = 1) is then solved for *i* = 1, …, *d*. Yata and Aoshima proposed a modification of eigenvalues [16] defined by:

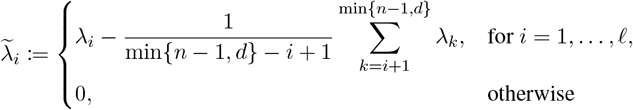

where *ℓ* ≤ min{*n* − 1, *d*} is a parameter representing the essential dimensionality. This modification improves the convergence of empirical eigenvalues to their population counterparts, indicating mitigation of the curse of dimensionality caused by noise accumulation. In the RECODE algorithm, the optimal value of the essential dimensionality *ℓ*^∗^ is automatically determined as

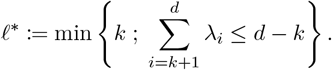

RECODE version 1 (RECODE v1) modifies the raw count matrix *X* such that the eigenvalues *λ*_*i*_ match the modified eigenvalues 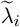:

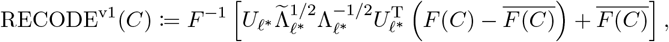

Where 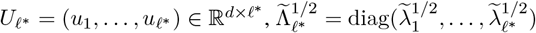, and 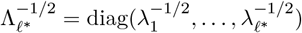.

Recently, Yata and Aoshima also proposed a correction method for eigenvectors [17]:

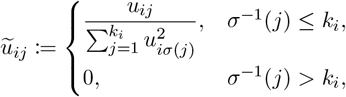

where *σ* : {1, …, *d*} → {1, …, *d*} is a permutation such that |*u*_*iσ*(1)_| ≥ · · · ≥ |*u*_*iσ*(*d*)_|, and *k*_*i*_ is the integer uniquely determined by the following condition:

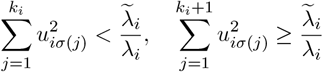

This eigenvector correction also alleviates the curse of dimensionality [17]. RECODE version 2 (RECODE v2) incorporates both the modified eigenvalues and eigenvectors:

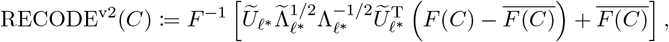

Where 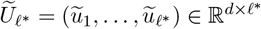.

### RECODE-based HVG selection

Let 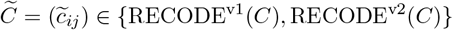 denote the scRNA-seq data denoised by RECODE. I apply two standard preprocessing steps, total-count normalization and log normalization, to the RECODE-denoised data 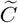. Given a non-negative matrix 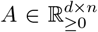, the total-count normalization *f* ^TN^ is defined as follows:

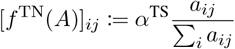

where *α*^TS^ is a scaling parameter known as the target sum or size factor. The log normalization *f* ^LN^ is defined as:

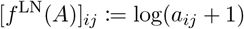

In RECODE-based HVG selection, I first apply total-count normalization followed by log normalization, resulting in the matrix 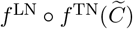. I then compute the variance 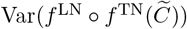, called *RECODE-denoised variance*, for each gene across cells and select the top n top genes genes with the highest variances as highly variable genes.

## Results

### Technical noise floor curve disturbs accurate HVG selection

To investigate the influence of technical noise on variability statistics, I first visualized the relationship between the mean expression per gene and several measures of variability under different normalization schemes (Figure 1a). In scRNA-seq data, a substantial number of genes, such as housekeeping or cell-cycle-related genes, are expected to be consistently expressed across most cell types and thus exhibit near-zero biological variance. Accordingly, variability measures should, in principle, include many values close to zero and remain largely independent of mean expression.

**Figure 1.**
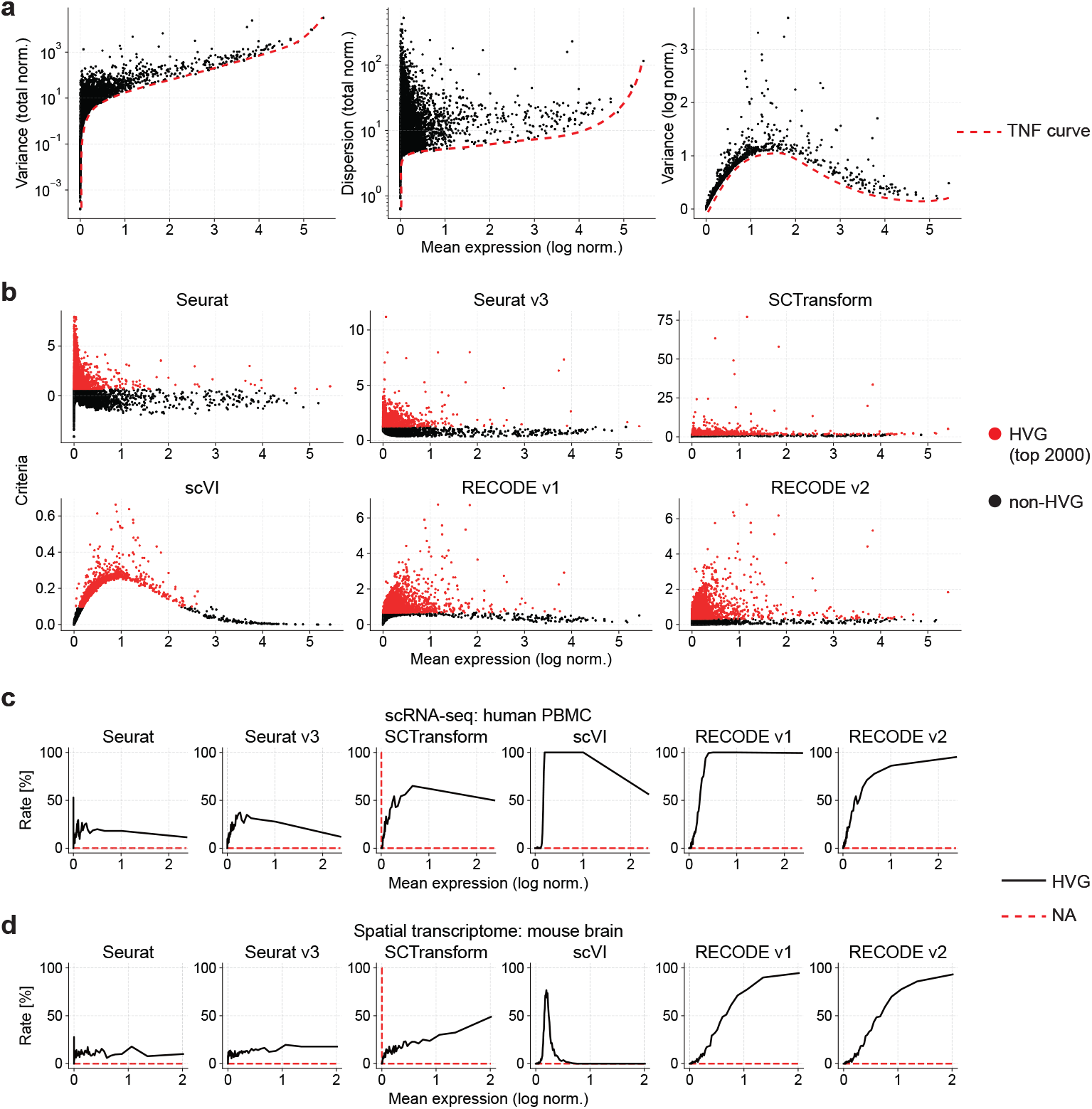
Overview of technical-noise structure and HVG selection. **a**, Scatter plots of mean expression per gene versus variability measures under different normalization schemes. Left: dispersion after total-count normalization; center: variance after total-count normalization; right: variance after log-normalization. Each black dot represents a gene, and the red dashed line indicates the technical noise floor (TNF) curve, representing the lower bound of variability statistics elevated by technical noise. **b**, Comparison of HVG selection criteria across six methods in human PBMC scRNA-seq data. The horizontal axis shows the criterion used by each method. Genes highlighted in red are those selected as HVGs with a cutoff of n top genes = 2,000, and genes in black represent the remaining genes. **c, d**, HVG detection rates (black lines) and missing-value (NA) rates (red dashed lines) as a function of mean expression, shown for the six methods in human PBMC scRNA-seq data (**c**) and mouse brain spatial transcriptomics data (**d**).

However, as shown by the red dashed lines in Figure 1a, the observed distributions of dispersion and variance after total-count normalization exhibit a pronounced lower bound that increases monotonically with mean expression. Even after log-normalization, the lower envelope of variance remains non-negligible, peaking around mean expression levels of 1–2 on the log scale. These lower bounds are not of biological origin but instead arise from technical noise intrinsic to the scRNA-seq measurement process. I refer to this increasing baseline as the *technical noise floor (TNF) curve*.

The existence of the TNF curve can be easily explained using a simple mathematical model. Let 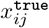 and *x*_*ij*_ denote the true and observed expression values, respectively, and let *e*_*ij*_ be the technical noise defined as their difference, 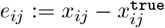. Assuming statistical independence between the true signal and noise, I obtain 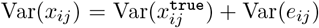. This relationship indicates that the observed variance Var(*x*_*ij*_) is elevated by the noise variance Var(*e*_*ij*_), regardless of the fluctuation of the true signal 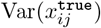. Therefore, the TNF curve mainly arises from the technical noise component *e*_*ij*_.

Genes whose expression variability lies near the TNF curve are likely dominated by technical noise rather than true biological variation, making HVG selection in these regions particularly error-prone. Therefore, reducing the technical noise that causes the TNF curve or equalizing the TNF curve across the range of expression levels is essential to enable fair comparison of gene-wise variances and to improve the accuracy of HVG selection.

### Different approaches have addressed the TNF curve in different ways

Next, I examined how effectively each existing method and the proposed RECODE approach account for the negative effects of technical noise, represented by the TNF curve, in detecting HVGs. For this analysis, I evaluated the distributions of each method’s criterion values and the expression ranges of the detected HVGs using the human PBMC scRNA-seq dataset (Figures 1b and 1c).

Because Seurat normalizes dispersion values within bins of mean expression using *z*-score normalization, the lower bound of normalized variance becomes irregular and fragmented across the entire range of mean expression. Seurat v3 corrects the variance by performing nonlinear regression based on a negative binomial model, resulting in an approximately horizontal lower bound. However, this correction tends to elevate the lower bound in the low-expression range, indicating that noise removal in this region remains incomplete. SCTransform modifies the target statistic from variance to dispersion and overcomes the tendency of Seurat v3 to detect many lowly expressed genes, yet several genes in the low-expression range remain unassigned (NA). Although scVI models gene expression through a hierarchical Bayesian framework, it shows limited improvement in reducing the TNF curve and mainly detects genes with moderate expression levels. RECODE v1 attenuates the TNF curve but does not completely flatten it, whereas RECODE v2 achieves an almost flat TNF curve approaching zero across all expression levels. This result indicates that technical noise can be effectively removed and that the variability statistics can primarily reflect biological differences. Importantly, this fundamental behavior was also demonstrated in simulated data in Imoto et al. (2022), where RECODE successfully reduced technical noise in Splatter-generated datasets and preserved the variance hierarchy between variable and non-variable genes, confirming that the mathematical foundation of RECODE-based HVG selection holds both in theory and in silico settings.

Furthermore, when compared with the mouse brain spatial transcriptomics dataset, which exhibits higher detection sensitivity (Figure 1d), the expression ranges of detected HVGs vary substantially among SCTransform, scVI, and RECODE v1. This suggests that the accuracy of these methods depends strongly on data quality and platform characteristics. In contrast, RECODE v2 yields highly consistent distributions between the PBMC and spatial transcriptomics datasets (Figures 1c and 1d). These results suggest that RE-CODE v2 can detect HVGs accurately and robustly, independent of data quality or experimental platform, which I quantitatively demonstrate in the following sections.

### Quantitative comparison shows high accuracy of RECODE-based HVG selection

In this section, I compared the accuracy of HVG selection using the same six methods as in the previous analysis (Seurat, Seurat v3, SCTransform, scVI, RECODE v1, and RECODE v2), applying them to both human PBMC scRNA-seq data and mouse brain spatial transcriptomics data. From the CellMarker 2.0 database, I selected 153 PBMC-associated marker genes and 102 mouse brain-associated marker genes for quantitative evaluation, and compared the HVG ranks assigned by each method based on their respective criteria (Figures 2a–2c).

**Figure 2.**
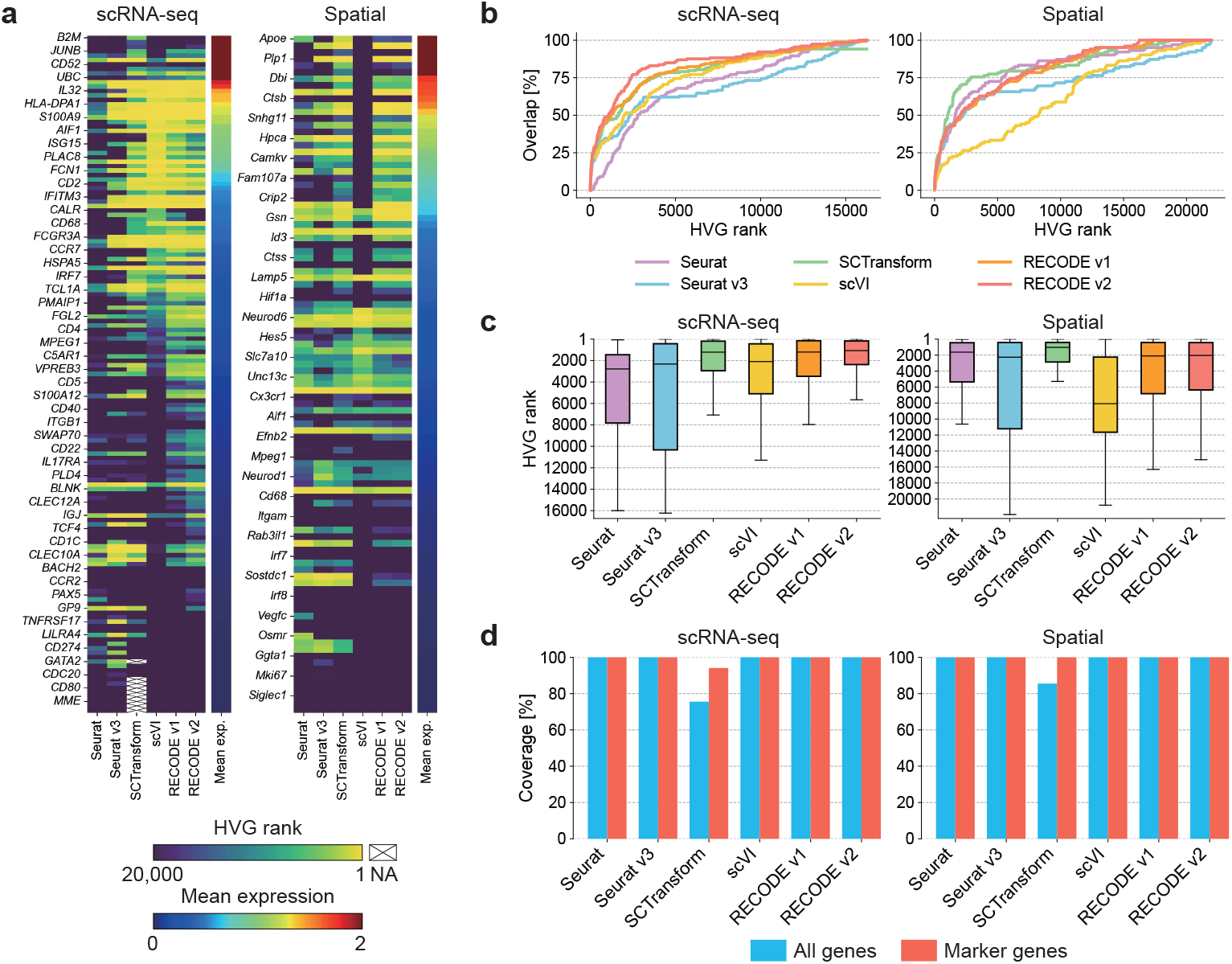
Comprehensive benchmarking of six HVG selection methods using scRNA-seq and spatial transcriptomics datasets. **a**, Heatmaps showing HVG rankings of known marker genes in the human PBMC scRNA-seq dataset (left) and the mouse brain spatial transcriptomics dataset (right). Genes are sorted by their mean expression levels. Boxes marked with a cross indicate missing values (NA). **b**, Overlap rates between HVG ranks and known marker genes as a function of the HVG rank for each method. **c**, Box plots of HVG rankings corresponding to panel **a. d**, Bar plots showing the coverage of ranked genes, defined as the proportion of marker genes (red) and all genes (blue) successfully assigned a rank.

When examining the dependence on gene expression levels, scVI detected marker genes primarily in the intermediate expression range, whereas the other methods showed no strong dependence on expression level (Figure 2a). In the accuracy comparison using known marker genes, SCTransform, RECODE v1, and RE-CODE v2 showed relatively high performance (Figures 2b–2c). Among them, RECODE v2 achieved the highest accuracy for the scRNA-seq dataset, while SCTransform performed best for the spatial transcriptomics dataset.

However, SCTransform failed to assign HVG ranks to a non-negligible number of genes, resulting in missing values (NA) (Figure 2d). This issue likely stems from the SCTransform procedure in which Pearson residuals are computed using regularized negative binomial regression; genes with extremely low expression or unstable residual variance cannot be robustly modeled, leading to missing HVG scores. Importantly, these missing values also occurred among marker genes, which is problematic for reliable HVG-based analyses.

In contrast, RECODE achieved both high sensitivity and complete coverage of marker genes in all datasets, confirming its robustness across different data types and measurement platforms. In the next section, I evaluate the robustness of these methods under simulated batch variation.

### Cell downsampling experiment highlights robustness of HVG selections

I next evaluated the robustness of HVG selection with respect to reductions in cell number. Using the same human PBMC scRNA-seq and mouse brain spatial transcriptomics datasets, I randomly downsampled cells to *n* = 2,500, 1,500, and 500, and performed HVG selection for each subset. For each *n*, this procedure was repeated 100 times, and I calculated the mean HVG rank of each marker gene together with the Jaccard similarity, which quantifies the consistency of detected HVGs across downsampled datasets (Figure 3).

**Figure 3.**
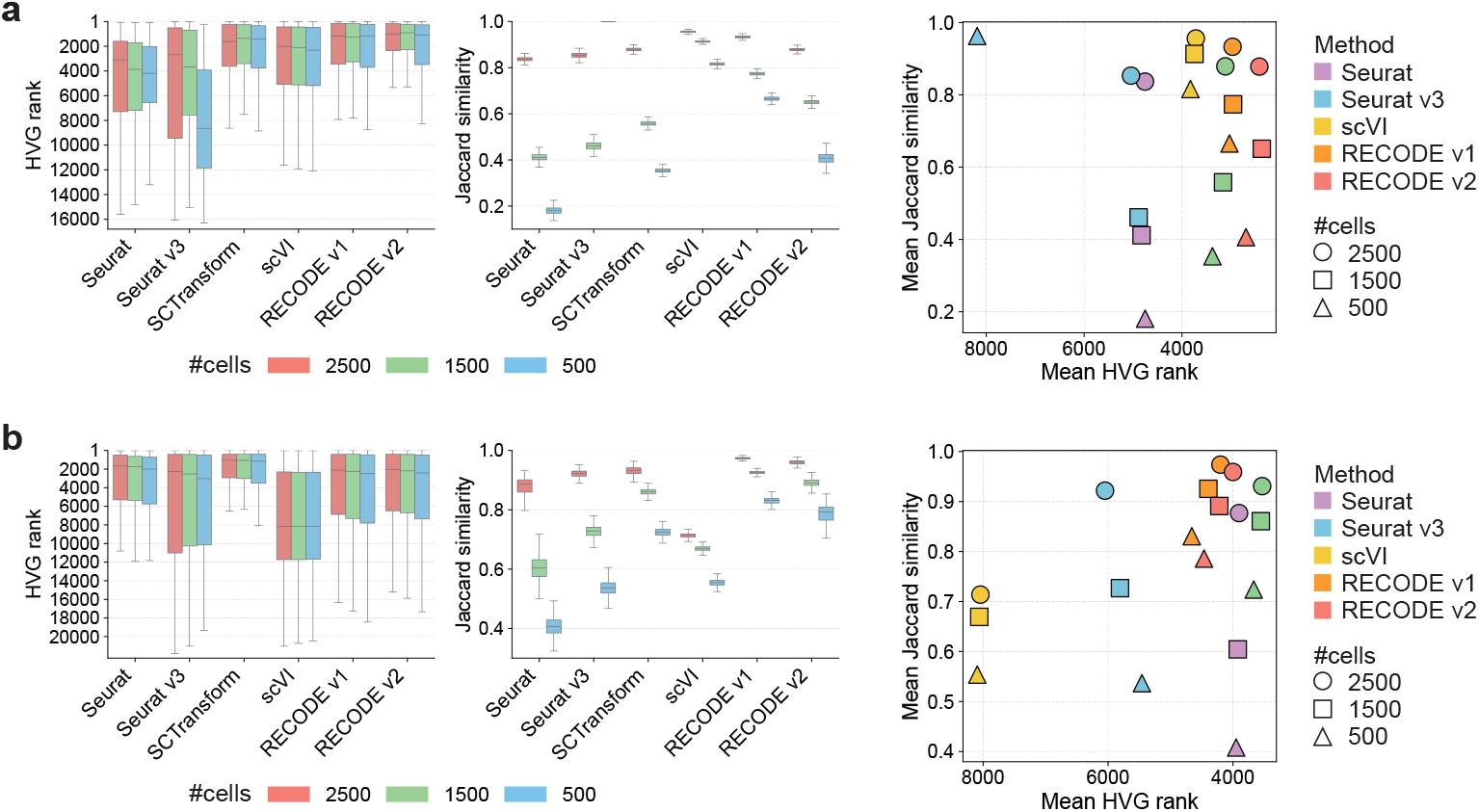
Robustness of HVG selection under cell downsampling. For the human PBMC scRNA-seq dataset (**a**) and the mouse brain spatial transcriptomics dataset (**b**), cells were randomly downsampled to *n* = 2,500, 1,500, and 500 for 100 iterations, and HVG rankings and Jaccard similarities were computed. Left: box plots of HVG ranks for known marker genes. Center: box plots of Jaccard similarity for the top 2,000 HVGs and scatter plots of the mean HVG rank versus the mean Jaccard similarity. A position toward the upper right indicates higher accuracy and greater robustness.

As expected, the accuracy of HVG ranking decreased as the number of cells was reduced (Figure 3a–b, left). This degradation was particularly severe for Seurat v3, which showed a marked loss of accuracy at smaller sample sizes. A similar trend was observed for robustness: the Jaccard similarity decreased with fewer cells, indicating less stable HVG selection under downsampling (Figures 3a and 3b, center). Although scVI was relatively robust for the scRNA-seq dataset, its performance deteriorated substantially for the spatial transcriptomics dataset, suggesting a strong dependence on data type.

Across both datasets, SCTransform, RECODE v1, and RECODE v2 demonstrated relatively high accuracy and robustness, with RECODE v1 and v2 showing slightly higher robustness than SCTransform (Figures 3a and 3b, right). In the next section, I investigate how these differences in HVG selection influence the accuracy of downstream analyses.

### Improved downstream clustering performance with denoised HVG-based representations

To examine whether the choice of HVGs and the use of denoised representations improve downstream clustering, I performed clustering analyses using the top 2,000 HVGs selected by each method. For SCTransform and the two RECODE variants, which perform denoising jointly with HVG selection, I used their recommended denoised data matrices. For all other methods, clustering was applied to log-transformed raw count data. Clustering was conducted using k-means, Leiden, and spectral clustering, which are well-used methods in single-cell biology. For each method, I varied random seeds and hyperparameters of the number of clusters or resolution to compute silhouette scores over 300 repetitions.

In the human PBMC scRNA-seq dataset, SCTransform, RECODE v1, and RECODE v2 consistently achieved higher silhouette scores across all clustering algorithms compared with Seurat, Seurat v3, and scVI (Figure 4a). These results indicate that HVG selection alone is insufficient for improving downstream clustering accuracy, and that denoising plays a crucial role in generating biologically meaningful representations.

**Figure 4.**
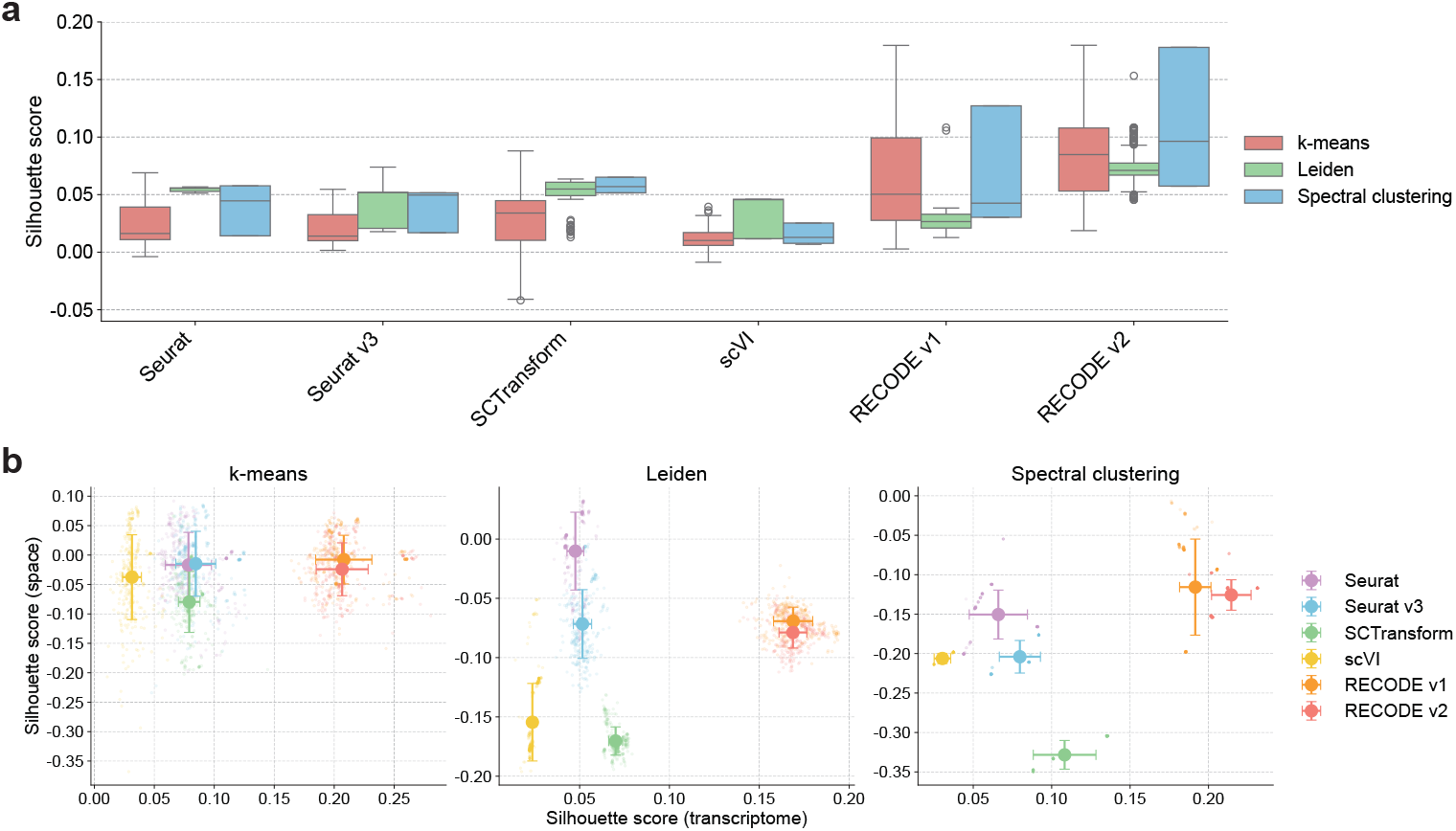
Quantitative evaluation of downstream clustering performance after dimensionality reduction with HVG selection. **a**, Silhouette scores of clustering on the human PBMC scRNA-seq dataset. Clustering was performed using k-means, Leiden, and spectral clustering methods, respectively, each repeated 300 times. Box plots summarize the distributions of silhouette scores across repetitions. **b**, Silhouette score distributions for the mouse brain spatial transcriptomics dataset under the same clustering conditions. The horizontal axis shows silhouette scores computed in the transcriptomic space, and the vertical axis shows silhouette scores computed in the spatial coordinate space. Large dots represent the mean scores, and error bars indicate standard deviations.

For the mouse brain spatial transcriptomics dataset, silhouette scores were computed in two coordinate systems: transcriptome space and physical spatial space. While SCTransform, RECODE v1, and RECODE v2 again produced relatively high silhouette scores in transcriptome space, SCTransform showed notably low silhouette scores when evaluated in spatial coordinates (Figure 4b). This suggests that although SCTransform effectively denoises transcriptomic variation, its clustering output does not align well with true spatial organization. In contrast, RECODE v1 and especially RECODE v2 maintained comparatively high silhouette scores in both transcriptome and spatial spaces, demonstrating that RECODE-based denoising supports clustering results that better reflect the underlying spatial structure of the tissue.

Together, those results demonstrate that the accuracy and robustness of HVG selection, as well as the precision of downstream clustering, are generally higher for RECODE v2 than for existing methods. In the next section, I discuss computational cost and summarize the overall findings of this study.

### Computational efficiency and overall performance summary

Finally, to evaluate the practical usability of each HVG selection method, I analyzed their computation time and scalability across different numbers of cells (Figure 5a). Scalability was defined as the logarithmic growth rate of computation time with respect to the number of cells, corresponding to the slope of the curves in Figure 5a. Because Seurat, Seurat v3, and scVI correct only variability statistics without modifying expression values, they achieved relatively fast runtimes. In contrast, SCTransform, RECODE v1, and RECODE v2 adjust both variance and expression levels through denoising or model-based normalization, resulting in slower overall computation. Among these three methods, however, RECODE v2 achieved the fastest runtime and demonstrated the best scalability. This efficiency arises because RECODE performs noise removal using only basic linear algebraic operations without requiring statistical parameter estimation of distributions or complex probabilistic models. Consequently, RECODE has inherently low computational cost for a denoising method, and RECODE v2 further optimizes the estimation of the noise-reduction function, enabling substantially faster execution relative to earlier versions.

**Figure 5.**
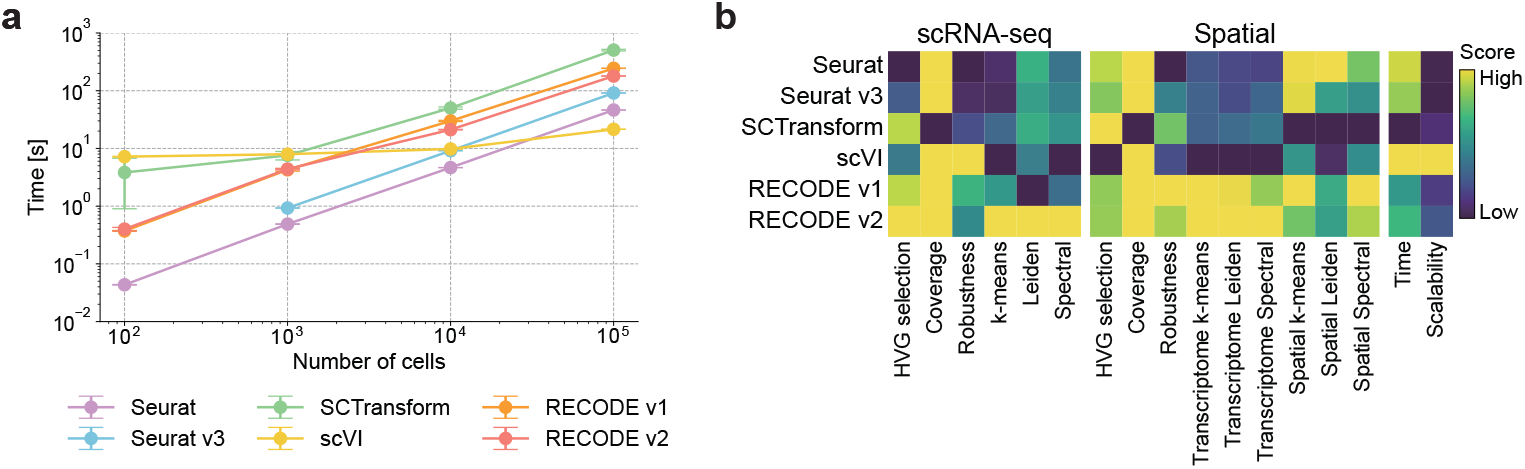
Computational cost and summary of overall performance. **a**, Computation time for the number of cells. For each cell count, the runtime was measured over 10 iterations; points indicate the mean, and error bars represent standard deviations. Seurat v3 failed at 10^2^ cells, and the corresponding data point is omitted from the plot. **b**, Summary heatmap of all evaluation metrics. Yellow indicates higher scores and dark blue indicates lower scores. Scalability is quantified as the regression slope of computation time between 10^3^ and 10^5^ cells, where lower values correspond to better scalability.

To integrate the results from all evaluation metrics, I summarized their normalized scores in a heatmap (Figure 5b). See the supplementary information for the method of computing scores. Although Seurat and Seurat v3 perform well in some individual metrics, their overall scores remain comparatively low, consistent with previous observations that they insufficiently model technical noise such as the TNF curve. scVI exhibits exceptionally low computational cost but faces persistent challenges in accuracy-based evaluations. The more recent methods, SCTransform, RECODE v1, and RECODE v2, achieved high scores across most categories, reflecting their mathematically principled noise modeling and effective removal of technical artifacts without compromising the true biological signal. However, SCTransform showed relatively low clustering scores in spatial coordinates, likely because it uses Pearson residuals from a negative binomial model as the representation for downstream analyses. Since these residuals lie on a scale different from true gene expression levels, they may not satisfy the assumption that transcriptionally similar cell types tend to occupy nearby spatial positions, thereby limiting performance in spatial-domain clustering.

Among all evaluated methods, RECODE v2 demonstrated the most balanced and superior performance overall. It achieved the highest score in 9 out of 17 evaluation categories (and 11 categories excluding RECODE v1) and simultaneously excelled in accuracy, robustness, downstream clustering quality, and computational efficiency. Together, these results position RECODE v2 as a high-performance and practically useful preprocessing method for both scRNA-seq and spatial transcriptomics analyses.

## Discussion

In this study, I performed a comprehensive evaluation of RECODE-based HVG selection, comparing it with widely used existing methods across both scRNA-seq and spatial transcriptomics datasets. The results showed that RECODE v2 achieves high accuracy in HVG selection, strong robustness under cell subsampling, reliable downstream clustering performance, and the best computational efficiency among denoising-based approaches. These findings demonstrate that RECODE v2 provides a well-balanced and high-performing solution for preprocessing single-cell transcriptomic data.

Across the accuracy-based evaluations, RECODE v2 consistently detected known marker genes with high performance and without missing values, whereas competing methods such as SCTransform frequently yielded NA values due to model instability. RECODE also exhibited stable HVG selection across datasets with distinct measurement characteristics, highlighting its robustness to variation in data quality and modality. Importantly, RECODE v2 achieved the best runtime and scalability among the denoising methods, benefiting from its reliance on linear algebraic transformations rather than statistical parameter estimation. Together, these results indicate that RECODE v2 effectively addresses the challenges introduced in the Introduction, namely, the presence of substantial technical noise, the distortion caused by the TNF curve, and the curse of dimensionality that affects downstream analyses.

Despite its advantages, RECODE has limitations. Because RECODE relies on mathematically precise modeling of technical noise, it assumes that the underlying noise structure follows the model implied by current single-cell sequencing technologies. If future sequencing platforms introduce noise characteristics that differ fundamentally from present-day technologies, the current mathematical formulation may no longer be optimal, and the algorithm may require modification to maintain accuracy. In addition, although this study did not evaluate multi-batch datasets, an extended version of the method, iRECODE [18], has been developed to simultaneously reduce both technical noise and batch-specific noise. This suggests that accurate HVG selection may be achievable even in multi-batch integration scenarios, an important direction for future validation.

Beyond methodological considerations, the results of this study provide broader implications for the field of single-cell analysis. The findings reinforce the view that noise removal at the raw data level is essential for reliable downstream analyses. Methods that correct only variability statistics or perform limited normalization cannot fully address the high-dimensional distortions introduced by technical noise. In contrast, RECODE’s mathematically grounded denoising restores biological structure without introducing artifacts such as cyclicity, enabling more trustworthy interpretations of clustering, gene expression variability, and cell-type identification.

In summary, the results of this study show that RECODE v2 is a high-performance denoising and HVG-selection framework that effectively resolves the fundamental noise-related challenges of single-cell transcriptomics. By improving accuracy, robustness, and computational efficiency, RECODE v2 serves as a reliable preprocessing solution for scRNA-seq and spatial transcriptomics, and it provides a strong foundation for future developments in single-cell data analysis.

## Data Availability

The scRNA-seq and spatial transcriptome datasets used in the comparisons of accuracy and robustness are the 3k Peripheral Blood Mononuclear Cells (PBMCs) from a Healthy Donor https://support.10xgenomics.com/single-cell-gene-expression/datasets/1.1.0/pbmc3k and Mouse Brain Section (Coronal) https://www.10xgenomics.com/datasets/mouse-brain-section-coronal-1-standard-1-1-0, respectively, provided by 10x Genomics. The scRNA-seq dataset used in the comparisons of computational time is the 1.3 Million Brain Cells from E18 Mice https://www.10xgenomics.com/jp/datasets/1-3-million-brain-cells-from-e-18-mice-2-standard-1-2-0 provided by 10x Genomics.

## Code Availability

The Python and R implementations of the RECODE algorithm, including functionality for highly variable gene selection, are publicly available at the following GitHub repository: https://github.com/yusuke-imoto-lab/RECODE

## Acknowledgment

This research was partially supported by the JST CREST (Grant Number JPMJCR24Q1).

## Author Contributions

Yusuke Imoto is the sole author of this study. He conceived the research idea, designed the methodology, conducted all analyses, and wrote the manuscript.

## Supplemental information

### Implementation of RECODE-based HVG selection

RECODE-based HVG selection is implemented in the `screcode` Python package (version 2.1 or later). Given an `AnnData` object [29] representing scRNA-seq data, users can perform HVG selection using the following procedure:

~~~
import screcode
recode = screcode . RECODE ()
adata = recode . fit_transform ( adata )
recode . highly_variable_genes ( adata )
~~~

By default, the top 2,000 genes with the highest variance are selected as HVGs. These genes are stored in `adata.var.RECODE_highly_variable`, and the RECODE-normalized variances used for ranking are stored in `adata.var.RECODE_normalized_variance`.

In addition, the RECODE-denoised data after applying total-count and log normalizations is stored in `adata.layers[“RECODE_log”]`. Users can use the following commands to set this as the default data matrix and subset it to HVGs:

~~~
adata . X = adata . layers[“RECODE_log”]
adata = adata [:, adata . var. RECODE_highly_variable ]
~~~

This procedure allows seamless reuse of RECODE-denoised data in downstream analyses using Scanpy or other libraries.

For R users, RECODE can be used by calling Python from within R using `reticulate`. For detailed instructions and tutorials, please refer to the GitHub repository at https://github.com/yusuke-imoto-lab/RECODE and the tutorial page at https://yusuke-imoto-lab.github.io/RECODE/Tutorials/index.html.

### Overview of the HVG selection methods used in this study

Here, I summarize the four highly variable gene (HVG) selection methods evaluated in the Supplementary Information and describe how each method was applied in this study.

#### Seurat (original method)

The Seurat HVG selection method [10] groups genes into bins according to their mean expression levels and then computes the dispersion (variance-to-mean ratio) for each gene. These dispersions are standardized within each bin using a *z*-score transformation, yielding normalized dispersion values (“dispersions norm”), which are used to prioritize HVGs. The method has been widely adopted due to its scalability and simplicity, and its practical performance has been validated in benchmark studies such as Yip et al. [9]. In this study, I applied Seurat’s method to log-transformed raw count matrices using the Python Scanpy implementation `scanpy.pp.highly_variable_genes(flavor=“seurat”)`. Although the original Seurat workflow includes a hyperparameter for filtering out lowly expressed genes, I did not use this filter to ensure a fair comparison among methods.

#### Seurat v3

Seurat version 3 (Seurat v3) [13] improved the earlier approach by introducing a variance-stabilizing transformation (VST), which corrects the observed mean–variance relationship using a negative binomial regression model [11]. The resulting normalized variances (“variances norm”) serve as HVG selection criteria. I computed Seurat v3 HVGs using the Scanpy implementation `scanpy.pp.highly_variable_genes(flavor=“seurat_v3”)` applied directly to raw count matrices.

#### SCTransform (version 2)

SCTransform extends the VST framework established in Seurat v3 to enable more robust normalization and multimodal data integration. Seurat v4 and v5 adopt sctransform version 2, which fits a regularized negative binomial regression model to stabilize variance across diverse datasets and modalities. In this study, I used the R `sctransform` package with `vst.flavor = “v2”` and employed the residual variance (“residual variance”) as the HVG selection criterion. The denoised Pearson residual matrix (“scale.data”) generated by SCTransform was used for downstream analyses such as clustering, in accordance with standard Seurat workflows.

#### scVI

scVI [14] is a deep generative model based on variational autoencoders that captures both biological variability and technical noise through a hierarchical Bayesian framework. For HVG selection, the scVI toolbox provides a Poisson-based gene selection routine, which ranks genes by quantifying their deviation from a Poisson model fitted to observed counts, effectively detecting genes enriched for biological variability relative to technical noise. I applied HVG selection using the `scvi.data.poisson_gene_selection` function in the `scvi-tools` Python library on raw count matrices.

### Marker gene selection

Marker genes used for evaluating HVG selection accuracy were obtained from *CellMarker 2* [30]. For the human PBMC scRNA-seq dataset, I first extracted immune cell-related marker genes and then selected only those corresponding to the cell types present in the dataset: *CD4 T cell, CD8 T cell, B cell, NK cell, Monocyte, Dendritic cell*, and *Platelet*. For the mouse brain spatial transcriptomics dataset, all marker genes annotated under the corresponding mouse brain-related categories in *CellMarker 2* were extracted without further filtering.

The complete list of extracted marker genes and the scripts used for selection are available on GitHub: https://github.com/yusuke-imoto-lab/RECODE_paper_materials.

### Clustering analysis

For the clustering analysis, I employed three widely used algorithms: *k*-means, Leiden, and spectral clustering, and evaluated their performance across multiple parameter settings. For *k*-means and spectral clustering, the number of clusters was set to 7, 9, and 11. These values were chosen to approximate the cluster numbers reported on the official 10x Genomics webpages for the corresponding datasets (9 clusters for both the human PBMC and mouse brain datasets).

The Leiden algorithm does not directly accept the number of clusters as an input; instead, I tuned the *resolution* parameter to values of 0.7, 0.9, and 1.1, chosen to yield cluster numbers comparable to those used in *k*-means and spectral clustering.

For each parameter setting, clustering was repeated 100 times with different random initializations, resulting in a total of 300 clustering runs per method. Clustering performance was quantified using silhouette scores computed from all 300 runs.

### Evaluation metrics

I quantitatively compared HVG selection methods using the following five evaluation metrics. These metrics assess accuracy, robustness, clustering performance, and computational efficiency.

#### HVG rank

For each method, HVGs were ranked according to the method-specific criterion (e.g., normalized dispersion for Seurat, normalized variance for Seurat v3, residual variance for SCTransform, Poisson-based statistics for scVI, and RECODE-denoised variance for RECODE). To evaluate accuracy, I examined the ranks assigned to known marker genes. Lower ranks indicate that a marker gene is prioritized more highly by the HVG selection method and thus reflects better performance.

#### Coverage

Coverage quantifies the proportion of genes for which an HVG score can be computed. This includes both (i) all genes and (ii) known marker genes. Coverage was computed as:

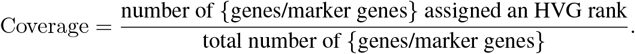

Methods that generate missing values (NA), such as SCTransform under low-expression conditions, exhibit reduced coverage.

#### Jaccard similarity

To evaluate the robustness of HVG selection under subsampling, I computed the Jaccard similarity between sets of selected HVGs obtained across repeated subsampling experiments. Given two HVG sets *A* and *B*, the Jaccard similarity is defined as:

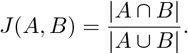

For each method, the top 2,000 HVGs were compared over 100 iterations at each subsampled cell count (*n* = 2,500, 1,500, 500), and the mean Jaccard similarity was used as a robustness metric.

#### Silhouette score

To assess downstream clustering performance, silhouette scores were computed using clustering results obtained from the top 2,000 HVGs. For scRNA-seq data, silhouette scores were computed in the transcriptomic feature space. For spatial transcriptomics data, silhouette scores were computed in two ways: (i) transcriptome-based and (ii) spatial-coordinate-based. The silhouette score for a cell *i* is defined as:

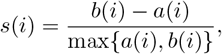

where *a*(*i*) is the mean intra-cluster distance for cell *i* and *b*(*i*) is the minimum mean distance between cell *i* and another cluster. Higher scores indicate better cluster separation.

#### Scalability

Scalability measures the computational efficiency of each method as the number of cells increases. For each method, computation time was recorded for datasets containing 10^3^, 10^4^, and 10^5^ cells. The scalability score was defined as the slope of the log–log regression line:

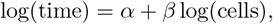

where a lower slope *β* indicates better scalability (i.e., slower growth in computation time as dataset size increases).

### Calculating scores

The scores shown in Figure 5b were computed using the evaluation metrics described in the previous section. Each metric was normalized so that higher values always indicate better performance, and all metrics were rescaled to the interval [0, 1]. For an evaluation metric *M* where larger values represent better performance (e.g., coverage, silhouette scores, Jaccard similarity), the normalized score was computed as:

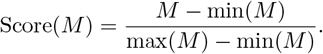

For metrics where smaller values indicate better performance (e.g., HVG rank, computation time, scalability), the metric was multiplied by −1 before applying the same normalization formula. Consequently, for every metric, the best-performing method receives a score of 1 and the worst-performing method receives a score of 0.

Figure 5b summarizes multiple performance dimensions using the following definitions:

- **HVG selection**: the mean HVG rank of known marker genes.
- **Coverage**: the coverage of all genes (proportion of genes assigned an HVG score).
- **Robustness**: the mean Jaccard similarity of the top 2,000 HVGs across 100 subsampling iterations.
- **k-means, Leiden, Spectral (Transcriptome/Spatial)**: the mean silhouette scores obtained from clustering using the respective algorithms.
- **Time**: the mean computation time for datasets containing 10^5^ cells.
- **Scalability**: the regression slope of computation time between 10^3^ and 10^5^ cells; lower slopes indicate better scalability.

This normalization procedure enables direct comparison across heterogeneous performance metrics and provides an integrated summary of the strengths and weaknesses of each HVG selection method.

## Supplemental figures

**Figure S1:**
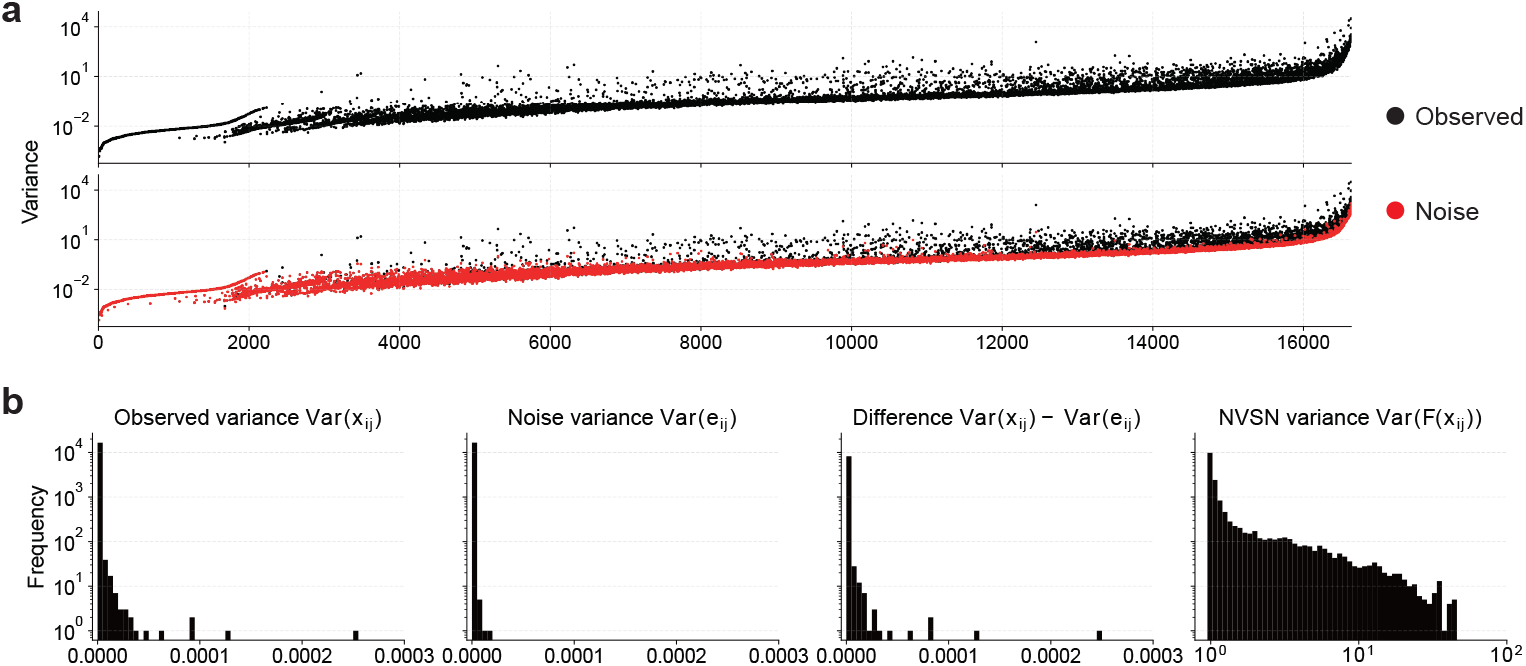
Relationship between observed variance and estimated noise variance. **a**, Scatter plot comparing the observed variance (black) and the estimated noise variance (red) for each gene. Genes are sorted from left to right by increasing mean expression. **b**, Histograms of the observed variance Var(*x*_*ij*_), estimated noise variance Var(*e*_*ij*_), their difference Var(*x*_*ij*_) − Var(*e*_*ij*_), and the variance after noise–variance stabilization (NVSN) Var(*F* (*x*_*ij*_)). The difference between observed variance and estimated noise variance is non-negative for all genes, Var(*x*_*ij*_) − Var(*e*_*ij*_) ≥ 0, confirming that the observed variance always exceeds the noise variance, as required by the basic variance decomposition property.

